# LISTER: Semi-automatic metadata extraction from annotated experiment documentation in eLabFTW

**DOI:** 10.1101/2023.02.20.529231

**Authors:** Fathoni A. Musyaffa, Kirsten Rapp, Holger Gohlke

## Abstract

The availability of scientific methods, code, and data is key for reproducing an experiment. Research data should be made available following the FAIR principle (**f**indable, **a**ccessible, **i**nteroperable, and **r**eusable). For that, the annotation of research data with metadata is central. However, existing research data management workflows often require that metadata should be created by the corresponding researchers, which takes effort and time. Here, we developed LISTER as a methodological and algorithmic solution to disentangle the creation of metadata from ontology alignment and extract metadata from annotated template-based experiment documentation using minimum effort. We focused on tailoring the integration between existing platforms by using eLabFTW as the electronic lab notebook and adopting the ISA (**i**nvestigation, **s**tudy, **a**ssay) model as the abstract data model framework; DSpace is used as a data cataloging platform. LISTER consists of three components: customized eLabFTW entries using specific hierarchies, templates, and tags; a ‘container’ concept in eLabFTW, making metadata of a particular container content extractable along with its underlying, related containers; a Python-based app to enable easy-to-use, semi-automated metadata extraction from eLabFTW entries. LISTER outputs metadata as machine-readable .json and human-readable .csv formats, and MM descriptions in .docx format that could be used in a thesis or manuscript. The metadata can be used as a basis to create or extend ontologies, which, when applied to the published research data, will significantly enhance its value due to a more complete and holistic understanding of the data, but might also enable scientists to identify new connections and insights in their field. We applied LISTER to the fields of computational biophysical chemistry as well as protein biochemistry and molecular biology, and our concept should be extendable to other life science areas.

## BACKGROUND

In a survey conducted in 2016 involving more than 1500 researchers, most respondents agreed that a reproducibility crisis exists in various scientific domains, including chemistry, biology, physics, engineering, and medicine^1^. Besides inherent factors related to the situation in academia, such as selective reporting, pressure to publish, poor analysis, and insufficient mentoring, extrinsic factors, such as the unavailability of methods, code, and primary data, i.e., data needed for reproducing an experiment, contribute to this crisis. The data unavailability led funding agencies and science publishers to require researchers to publish their data. For example, the German Research Foundation (DFG) expects research data to be made available for at least ten years^2^.

Conscious efforts to build better data infrastructure and standards have received wide attention. E.g., the National Research Data Infrastructure (NFDI) was established in Germany as an association comprised of many German institutions whose purpose is to create a permanent digital knowledge repository. Almost 20 NFDI consortia have formed in the natural sciences, life sciences, engineering, humanities, social sciences, and cultural sciences. Soon, up to 30 consortia will be funded by German federal and state governments^3^. In the US, the National Science Foundation (NSF) requires research data obtained under the NSF grants to be shared and to adhere to guidelines specifically designated for each NSF directorate field^4^. As a publisher, the American Chemical Society (ACS) also provides a guideline regarding the data policy: Each journal under the ACS umbrella follows one of four policy levels concerning data availability^5^.

Research data needs to be made available following the FAIR data principle, which provides guidelines as to four main quality aspects: Research data should be **f**indable, **a**ccessible, **i**nteroperable, and **r**eusable^6^. Further specifications have been recommended for dedicated research fields. E.g., for machine learning, Heil et al. categorize three research data reproducibility standards based on which data quality factors are satisfied^7^.

Besides the question of reproducibility, one can envision several use cases that can benefit from good research data management, ranging from influences on the local research environment to global scales. I) Data preparation and collection can be made easier for researchers before, e.g., thesis, article, or grant proposal preparation when regular Research Data Management (RDM) practices are applied. II) An RDM system allows for finding and filtering research data produced by fellow lab members or ex-lab members, e.g., for extending the research series. III) Data entries in a data catalog and associated with metadata, i.e., highly structured data documentation, can be aligned to other data repositories in different levels of granularity and technology stacks.

We reasoned that several requirements need to be satisfied to support such use cases:

1. Primary data should be accessible and downloadable openly when the license is not restrictive (which is mostly the case in scientific publications).
2. The metadata of these primary data should be extracted using minimum efforts to allow researchers to focus on the experiment design, execution, and analysis instead of manually extracting the metadata.
3. Whenever possible, an existing and proven technology stack or standard should be adopted.
4. Both primary data and metadata should be stored on a long-term basis with regular maintenance and backup operations.

The annotation of (primary) research data with metadata is central to any RDM workflow. Some specifications and frameworks exist to standardize metadata, both structurally and terminologically. As one, the ISA Model provides a community-driven framework for managing heterogeneous experiments in the life science, biomedical and environmental domains^8^. The model is based on three fundamental concepts surrounding *Investigation* to describe the project context, *Study* to explain the unit of research, and *Assay* to provide analytical measurements^8^.

Designing an RDM workflow requires considering how the data are collected, which includes the use of lab notebooks. Traditionally, lab notebooks have been written on paper. However, this is deemed impractical due to the difficulties, e.g., with archival and searchability of the entries. Electronic lab notebooks (ELNs) offer an additional set of features and have been adopted in many research groups. We focus on eLabFTW^9^, a web-based, open-source electronic lab notebook software that can be installed on a server and features common ELN functionalities such as experiment tracking, lab asset management, timestamping (as legal proof in the case of patent disputes), lab equipment scheduler, experiment documentation, and the possibility to create custom-type entries as database^10^. eLabFTW facilitates metadata storage that can be attached to each experiment and is widely used in the scientific community (in GitHub, it has > 700 stars, and > 1700 issues have been submitted, with ~1600 issues resolved/closed)^11^. eLabFTW is freely available for download and use and is currently being translated into 17 languages.

## SCOPE

Here, we provide methodological and algorithmic solutions to satisfy the first and second requirements mentioned above, focusing on the field of life sciences, which includes molecular modeling/simulations and wet lab experiments as pilot cases. To this end, we: I) customize eLabFTW entries using specific hierarchies, templates, and tags, II) provide a ‘container’ concept in eLabFTW, making metadata of a particular container content extractable along with its underlying, related containers, and III) develop an app to enable easy-to-use, semi-automated metadata extraction from eLabFTW entries, along with providing a workflow to support this extraction adopting existing and proven technology standards; the latter satisfies the third requirement. We coined this approach LISTER (**Li**fe **S**cience Experimen**t**s Metadata Pars**er**).

Our developments are guided by the necessities of users whose scientific focus should not get distracted by manually annotating experiments to conform with RDM and, in particular, metadata creation and extraction. This aspect makes LISTER unique compared to alternative approaches^12^.

## APPROACH

In this section, we elaborate on the requirements from above and the designed workflow. As a design principle, we used existing RDM standards and platforms whenever possible (see below). That way, we can focus on tailoring integration between existing platforms and developing necessary applications as a critical contribution to implementing good RDM practices. We use eLabFTW as the ELN and adopt the ISA model as the abstract data model framework, which is implicitly supported by our eLabFTW adaptation. DSpace, an open-source repository software package typically used for creating open-access repositories for scholarly and/or published digital content^13,14^, is used as a data cataloging platform.

### Design

In Figure 1, the design of the RDM workflow based on the requirements is illustrated. Steps (a) – (f) of the workflow, including the respective detailed steps, will be introduced below.

a. *Incorporating the ISA data model into eLabFTW*. As the ISA model is an abstraction of three main levels of research activities (Investigation, Study, Assay) and the relationship among those levels, a more concrete adaptation of the ISA model is required for application within eLabFTW. Our adaptation provides a template for life science-centered research, although adaptation details might vary for different research groups, depending on the perceived meaning and granularity of the activity levels within the group. In addition, metadata regarding a project (e.g., project title, responsible person for the project) and additional information (e.g., as to a publication) is provided through an extension of this model within our eLabFTW implementation. This will be described in more detail below.

i. *Mapping of the ISA data model to eLabFTW*. The native eLabFTW ‘class’ types contain *Experiment* and *Database*. ISA data model’s *Assay*, by definition, can be mapped directly to eLabFTW’s *Experiment*, which contains the experiment documentation. To map ISA’s *Investigation* and *Study*, for which no direct equivalents are available in eLabFTW, we use eLabFTW’s *Database* type, as it is a customizable abstract class from which a specific class implementation can be created by inheritance. The inherited class has a template that requires the user to enter pre-specified information. We map ISA’s *Investigation* to the specific class *System*, which represents, e.g., the central molecular target or hypothesis investigated. ISA’s *Study* is mapped to the homonymous specific class in eLabFTW, which provides context regarding the study subject and characteristics and groups *Experiment* instances. The relationship between these three basic classes is shown in Figure 2a. A *System* entry contains further information, such as the system name and the responsible person (Figure 2b). A *Study* entry contains information on the study’s aim and the responsible person, along with the study’s start and end date (Figure 2b). A *System* entry is a container for multiple *Study* and *Experiment* entries, and a *Study* entry is a container for multiple *Experiment* entries; the “has a” associations are realized via links in eLabFTW (Figure 2b). That way, questions can be asked such as “which studies were performed for a system” or “which experiments belong to a study or a system”. Additionally, metadata to be created for an *Experiment* entry can be augmented by metadata from the *Study* and *System* containers. We intentionally keep redundant associations between containers/entries. For example, while there is an indirect association between *Experiment* and *Project* via *Study* and *System* (see Figure 2b), we still keep the direct association between *Experiment* and *Project*, as this helps users of eLabFTW to identify the *Project* to which an *Experiment* belongs without having to go through the *Study* and *System* entries from the *Experiment*. Additionally, it disambiguates the *Project* corresponding to the *Experiment*, since the cardinality between *Project* and *System* is `m:n`.
ii. *Extending the ISA Model within eLabFTW*. We created three other classes (*Project, Publication*, as well as *Protocol/Materials and Methods* (*MM*)) in the eLabFTW hierarchy from the abstract class *Database* (Figure 2b). A *Project* complements the *System* class, and a *System* entry can have one or multiple *Project* entries associated. As the name implies, this class contains attributes related to a project, such as cooperation partners or the project manager. A *publication* is a container for experiments that have been reported in a publication. That way, metadata and primary data associated with that publication can be directly extracted, e.g., for submission to the publisher. *Protocol/MM* serves as a library of annotated template protocols/MMs (see the chapter *Generating reusable experiment documentation* for details), from which *Experiment* entries will be derived. The complete metadata entry list for *System, Project, Study*, and *Publication* container types is available in SI Chapter 2, Table S1. In addition to linking experiments/class entries with other class entries/experiments in eLabFTW, we are using tags to organize entries to make them better findable by using the filtering mechanism provided in eLabFTW. Exemplary tags are shown in Figure 3, organized into three main categories: *lab type tags* (e.g., dry lab *versus* wet lab), *system tags*, and *method tags*. Creating these tags requires domain expertise and an overview of the lab organization. Other categories than the three introduced here may be relevant for other labs.
b. *Generating reusable experiment documentation*. Research groups typically have cataloged documentation of how specific experiments are performed. We use such documentation, eventually redesigning it to become broadly reusable templates. We distinguish two categories of experiment documentation: protocols versus MM descriptions. Protocols, also termed standard operating procedures (SOP), are step-by-step guidelines for how to conduct a specific experiment and can be typically very detailed. MMs provide a textual description typically found in journal articles; MMs can often be generated by condensing the protocols. In our case, the MM descriptions are used to derive experiment documentation in eLabFTW, from which MM sections for theses or paper manuscripts can be automatically generated. The graining of protocols and MMs should reflect the modularity of experimental procedures used in a lab without creating too much overlap or redundancy. While this process may take some time and require expertise, the templates can be reused efficiently when a related experiment is done. This saves scientists from writing the protocols or MMs from scratch to document their experiments, although likely experimental details need to be adapted. Experiment documentation can be derived from a template stored in the *Database*-inherited class *Protocols/MMs* by importing the template content using functionality available within eLabFTW: it is triggered by the hashtag symbol followed by the words used as the title in the referred database entry. For changing or revising the protocols and MMs, we exploit that eLabFTW saves the revision history; hence, the revision history can be referred to using a permanent link pointing to the version that has been used to derive an experiment.
c. *Annotating reusable experiment documentation for automated metadata extraction*. To allow automated extraction of metadata (e.g., key-value pairs, which can be augmented with measure and unit information; see Figure 4), the protocol and MM templates are annotated with markups of the LISTER annotation language. The key-value pairs can be extracted with the LISTER extraction app (see Figure 5 for the overall interplay between these components). Metadata generation is necessary before providing primary data for archival in institutional, publisher, or community-wide repositories to fulfill the FAIR principles. The combination of annotated reusable experiment documentation and automated metadata extraction alleviates the time-consuming, cumbersome, and error-prone hurdle of annotating primary research data with metadata by hand each time such primary data is generated. Alternatively, metadata generation may be taken over from research equipment output, e.g., log files of a microscope and recording details in connection with microscopy data, which later may be augmented with additional metadata^15^.

i. *Annotating reusable experiment documentation*. Domain experts in the respective research fields or working groups will add markups to the protocol or MM templates adhering to the format of the LISTER annotation language. A brief example of an annotated MM is given in Figure 4; more detailed descriptions are given in the Implementation section as well as in SI Chapters 1.3 and 4. The design of the LISTER annotation language was guided by simplicity principles, letting researchers use the implementation without having a steep learning curve. The LISTER annotation preserves the readability of the text and, that way, mimics other markup languages, such as Markdown or the long-known HTML style (but with fewer annotation symbols striving for more readability). It allows including comments, iterations, and conditionals. Some additional principles for writing MMs are described in SI Chapter 3.
ii. *Adapting parameters and extracting metadata from annotated experiment documentation*. After scientists import the annotated protocol or MM templates relevant to their experiment from the *Protocols/MM* class into their *Experiment* entry, they likely need to adapt the predefined parameters according to the experiment details. Additionally, irrelevant parts of the templates should be removed, and new parts can be added, complying with the format of the LISTER annotation language. The LISTER app checks the syntax of the annotation markups, e.g., with respect to unmatched brackets or the number and types of elements in a key-value pair, as a validation mechanism upon parsing the experiment entries. This will yield either a warning message (when the issue does not affect the validity of the output) or an error message (when the issue affects the validity of the output), pointing to the problematic line(s) in the evaluated experiment entry (see SI Chapter 1.5). After the check, metadata is extracted from the annotated experiment documentation with the app (Figure 4a).
iii. *Parsing a group of experiments under the same container class. Experiment* entries can be parsed for metadata individually via LISTER’s GUI (Figure 6). Additionally, another important use case is to parse the content of a container, such as a *Publication* entry. A publication can be linked to several *Experiment* entries, which were conducted for the publication, and implicitly to *System* entries, *Project* entries, and *Study* entries (Figure 7). In the Database tab in LISTER’s GUI (Figure 6), the eLabFTW ID of a *Publication* entry can be provided, and LISTER will extract the metadata output from all experiment documentation under that publication, as well as from the associated *System, Project*, and *Study* entries.
d. *Using well-known input and output formats*. As input, LISTER parses the content of annotated, adapted experiment documentation in eLabFTW provided as HTML pages via the eLabFTW API (Figure 4a). LISTER transforms the content of annotated documentation into several outputs (Figure 4b, c): I) Experiment documentation as “clean” text without annotations to be used as MM sections in theses and manuscripts is provided in .docx format; II) Contextual experiment metadata is provided in .xlsx and .json formats, the first as human-readable version and the second as machine-readable version, e.g., to be used as input for DSpace’s data cataloging platform (Figure 4d).

**Figure 1.**
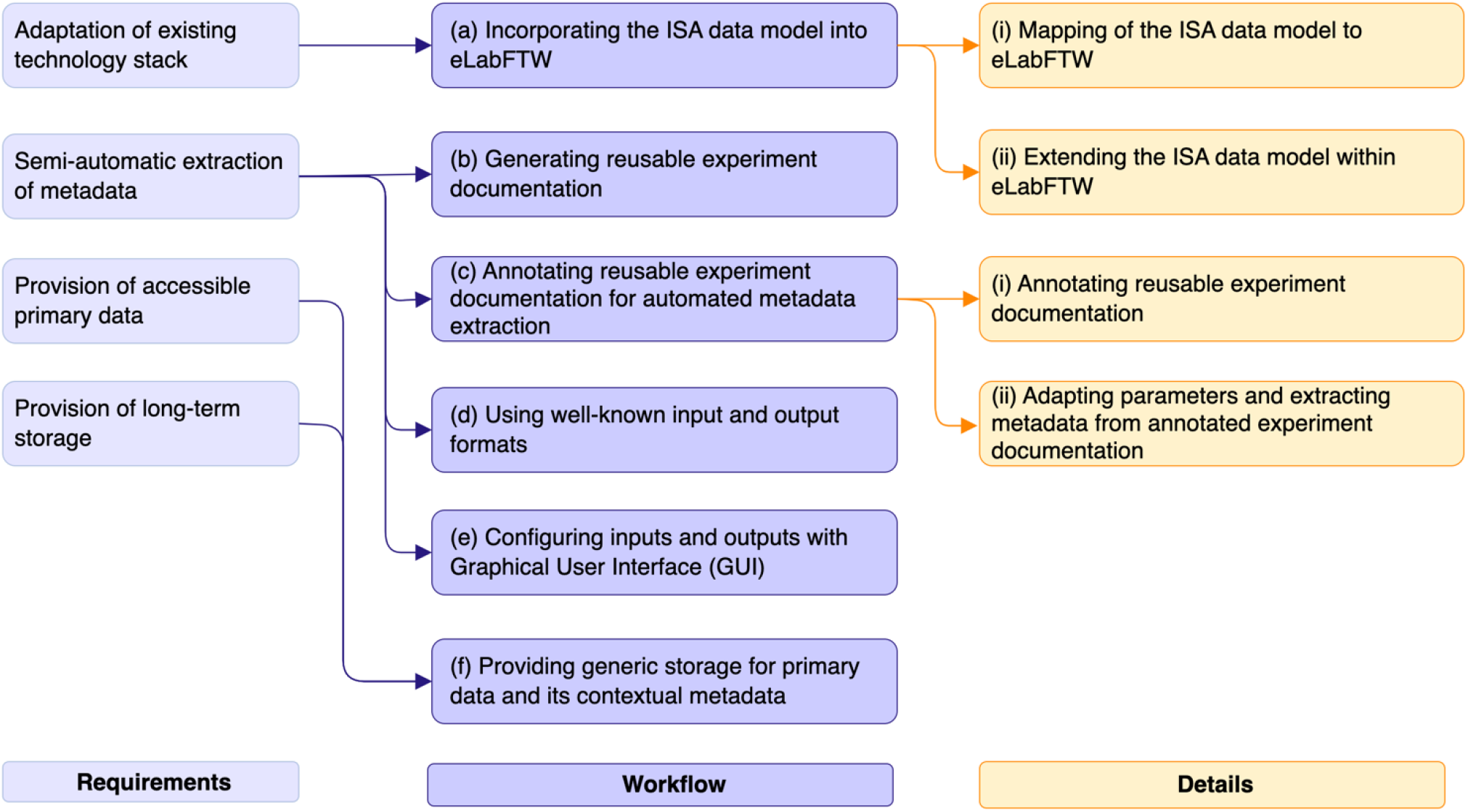
Requirements (left) and developed workflow (middle) to make primary data accessible and downloadable and extract metadata using minimum effort. Detailed steps are given on the right.

**Figure 2.**
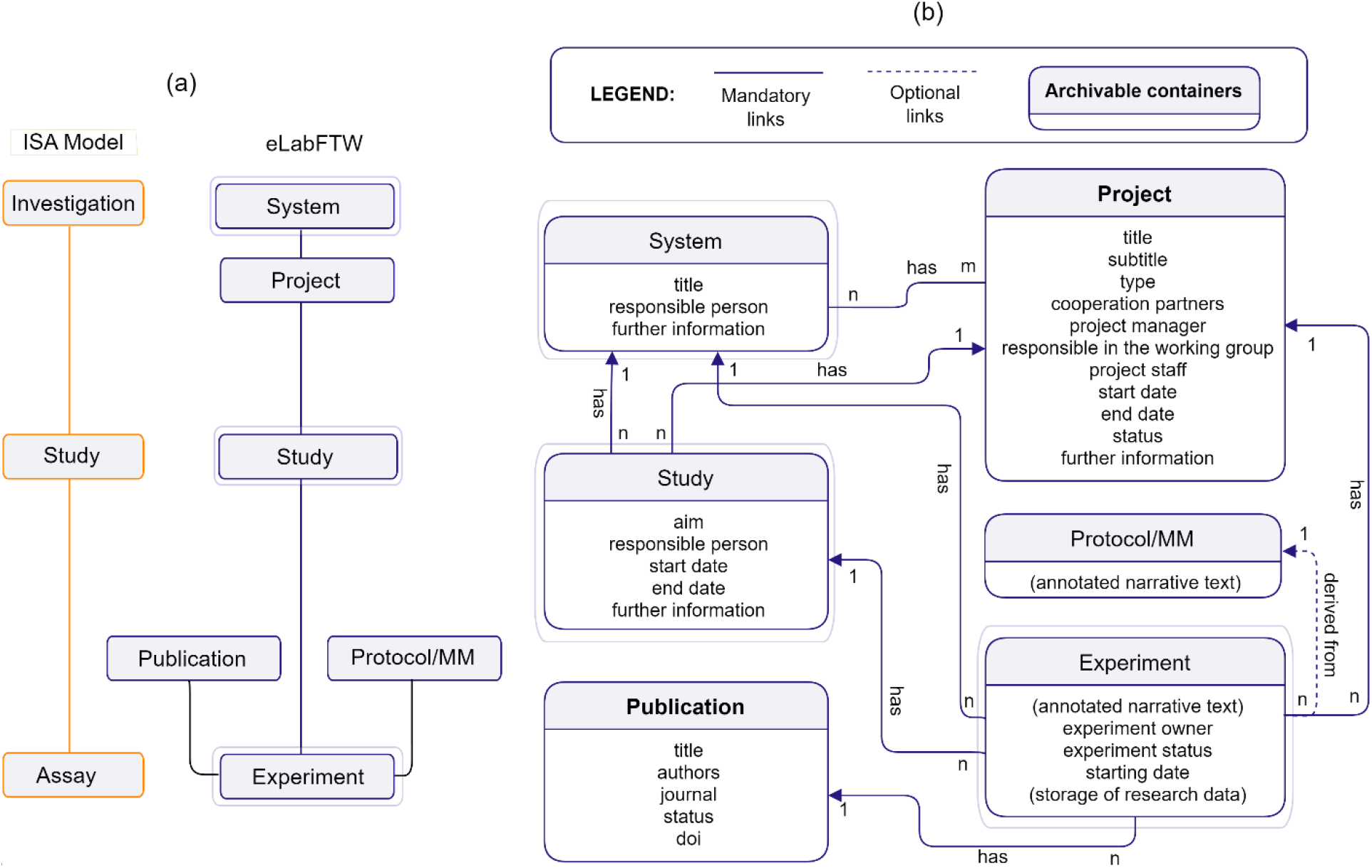
Incorporation of the ISA data model into the eLabFTW hierarchy of ‘class’ types. (a) Mapping between the ISA model (orange boxes) and eLabFTW classes (boxes with gray outer line); for *System* and *Study*, specific classes were inherited from the *Database* class in eLabFTW. In addition, specific classes *Project, Publication*, and *Protocol/MM* were inherited from *Database*. (b) Attributes of and relationships between specific class entries in eLabFTW. *Protocol/MM* and *Experiment* entries have no fixed attribute fields. Solid arrows indicate mandatory relationships, dashed arrows optional ones; “has” denotes an association, where the arrowhead points to the container.

**Figure 3.**
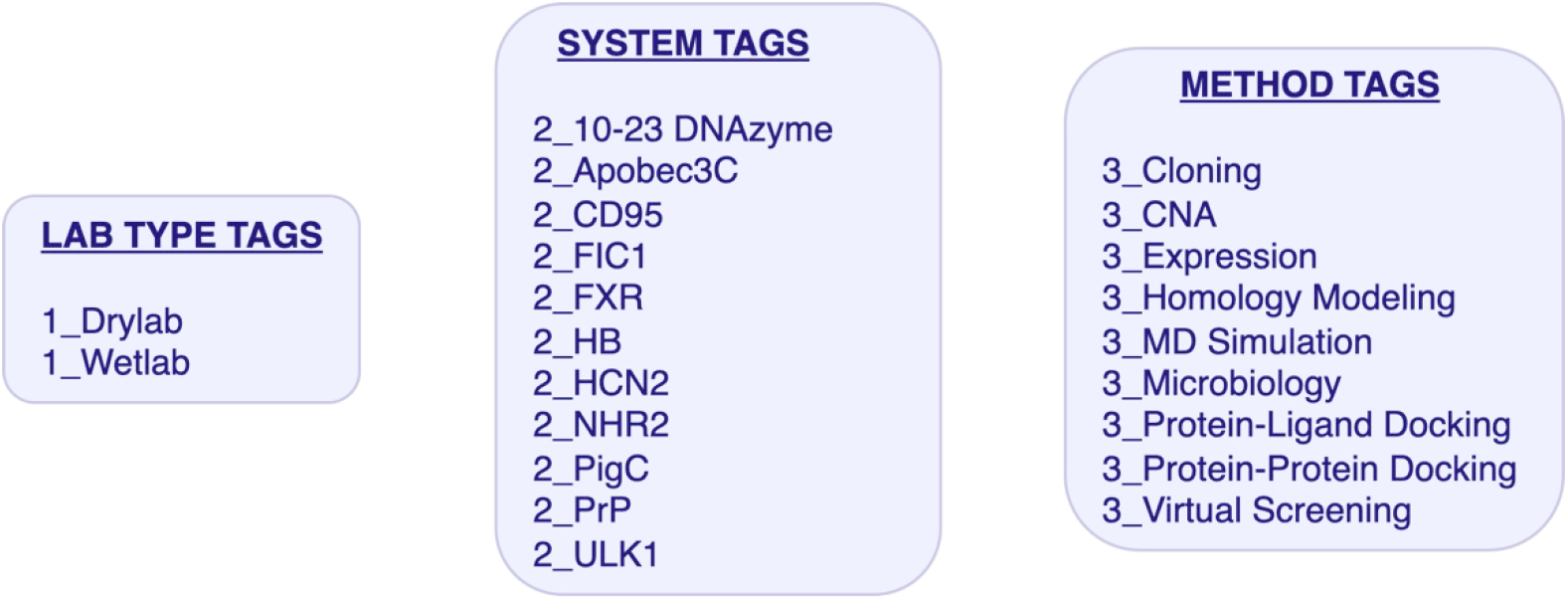
An exemplary implementation of a tag structure. Each experiment or relevant class entry (e.g., protocols or MM) should contain tags, which are used to categorize the entries and make them better findable using the filtering mechanism provided in eLabFTW.

**Figure 4.**
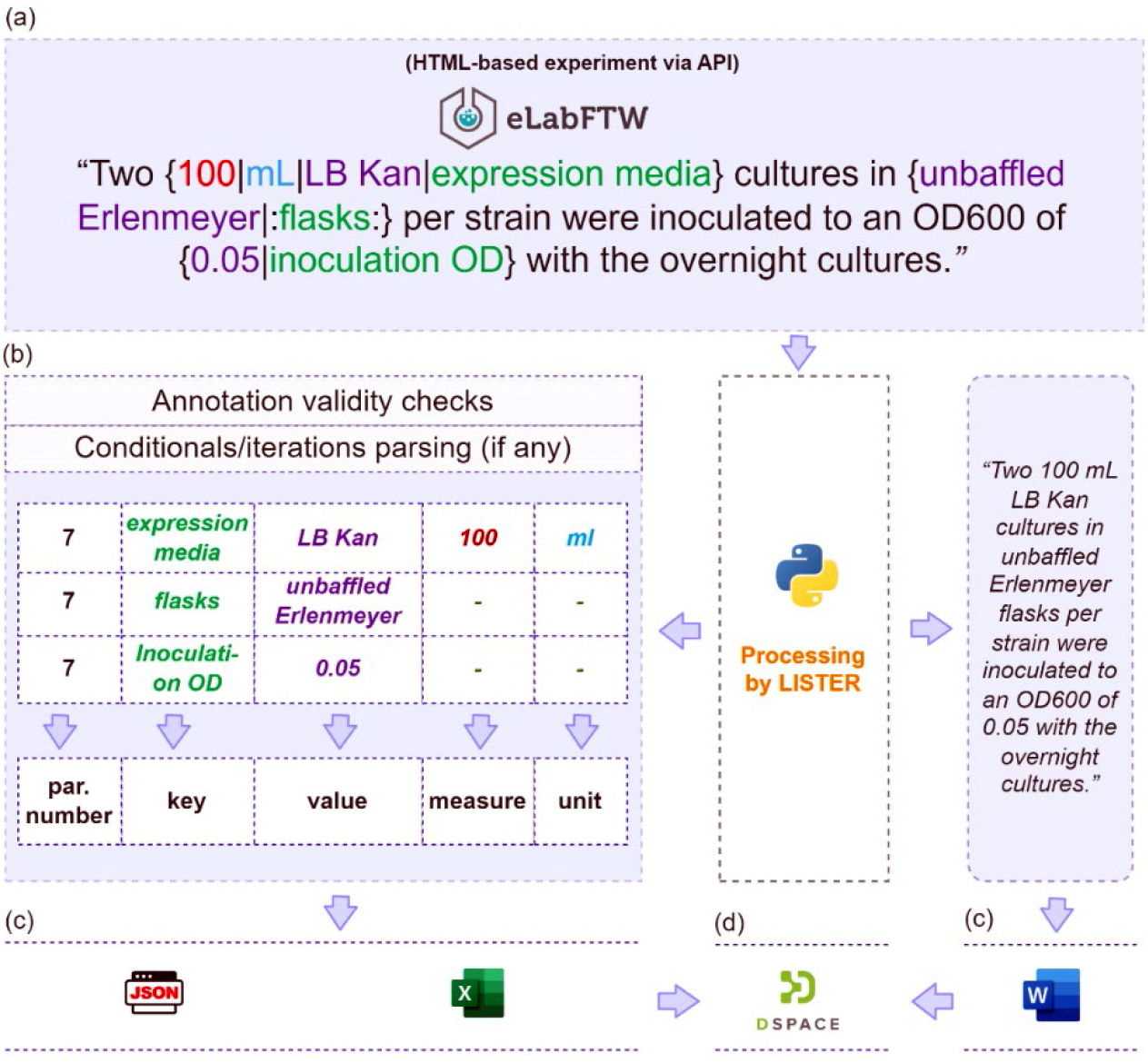
LISTER annotation language and results of metadata extraction with the LISTER extraction app. (a) Annotated and adapted experiment documentation is provided as an HTML page via the eLabFTW API as input to the LISTER app. (b) Each LISTER annotation markup is transformed with the app into a section number and corresponding key-value pairs along with, if available, measure and unit information. (c) LISTER’s output comprises metadata available in .xlsx and .json formats as well as clean, readable text in .docx format. (d) The metadata together with the primary data can be stored, e.g., in the DSpace catalog.

**Figure 5.**
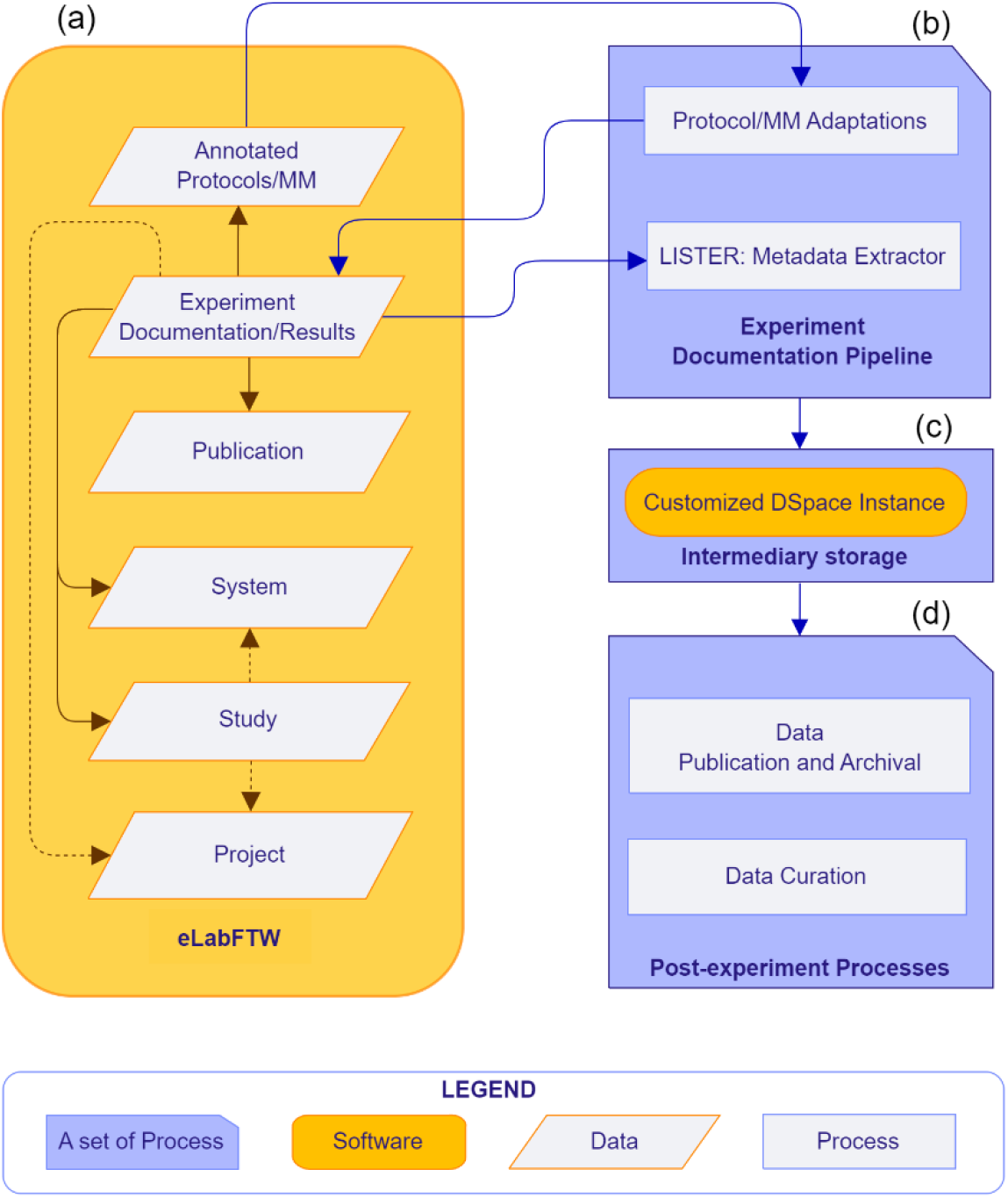
The interplay of the components of the LISTER-based RDM workflow. (a) eLabFTW contains a hierarchy of specific *Database* classes as containers to map an (extended) ISA model, stores annotated protocol and MM templates; and holds adapted annotated experiment (Figure 4a) documentation and results. (b) The user adapts annotated protocols and MMs and invokes metadata extraction with LISTER via the GUI, as seen in Figure 4b. (c) Extracted metadata and corresponding primary data are stored in, e.g., a customized DSpace instance as intermediary storage (Figure 4d). (d) From the intermediary storage, metadata and primary data can be prepared for archival (e.g., in Zenodo and bioRxiv) and curation (e.g., with Re3Data) or retrieved for publication purposes (e.g., on publisher websites).

**Figure 6.**
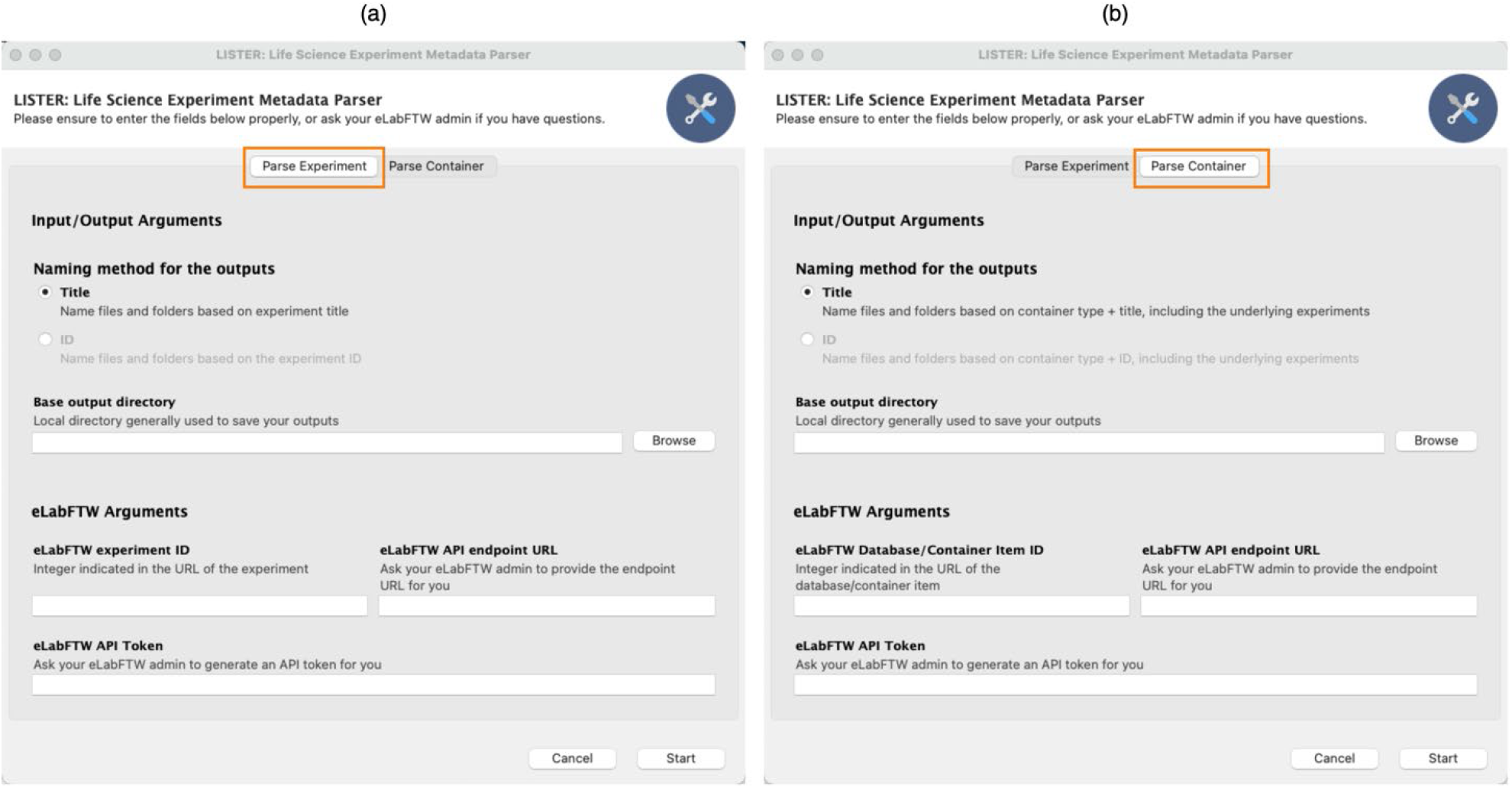
Graphical user interface (GUI) for LISTER. The app has two interface tabs: (a) Experiment – in which users can extract metadata from an *Experiment* entry, and (b) Container – in which users can extract metadata from a specific type of ‘container’, e.g., a *Publication* entry. In the latter case, the metadata from a *System, Project*, or *Study* of the *Experiments* that are linked directly to the *Publication* will also be extracted.

**Figure 7.**
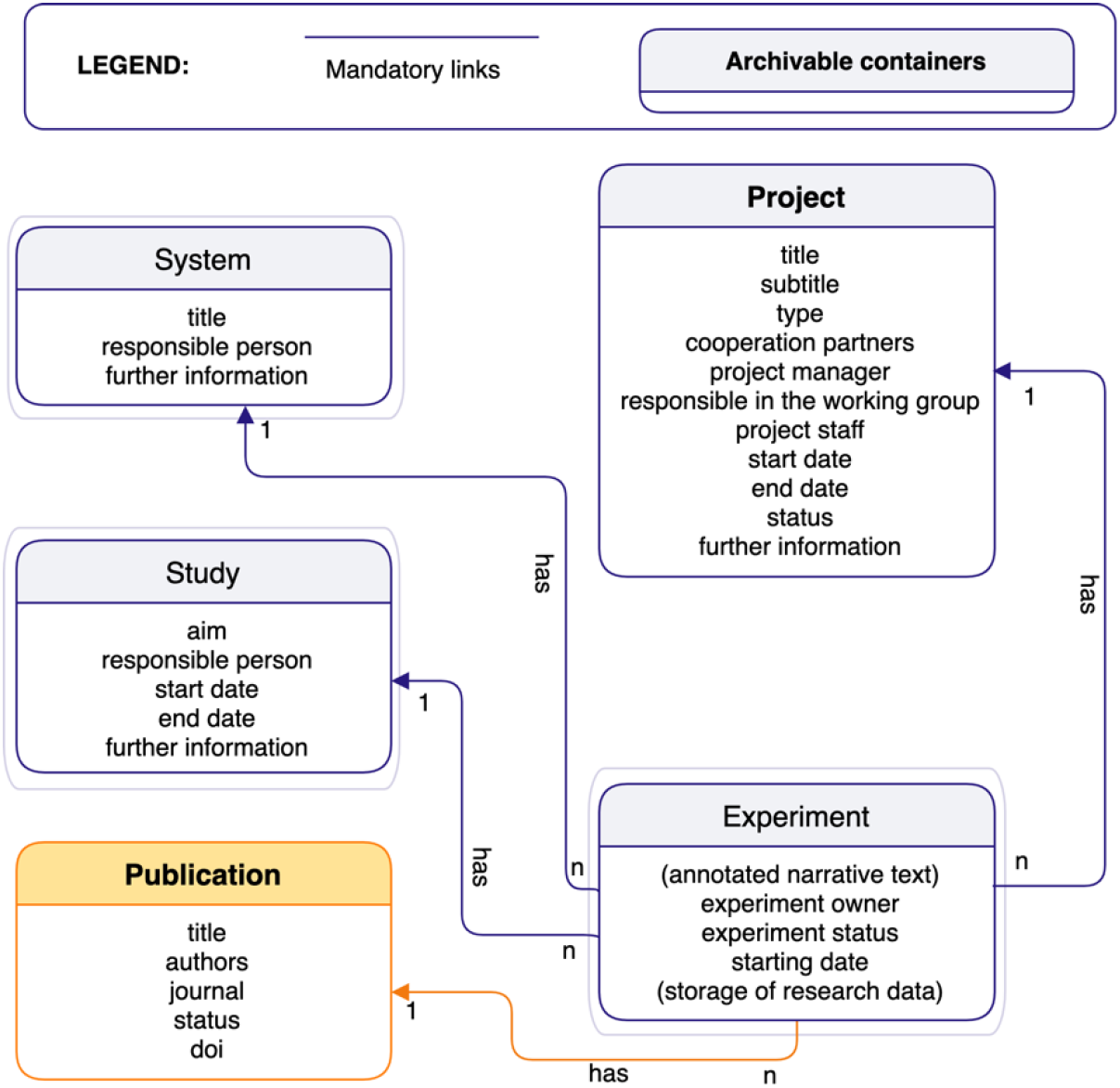
Illustration for a *Publication* entry (orange rectangle) acting as a ‘container’ from which metadata is extracted with the LISTER app. LISTER also extracts metadata from the corresponding *System, Project*, and *Study* that is directly linked to the relevant experiments concerning the extracted *Publication*.

### Methods

#### LISTER annotation language

We briefly summarize LISTER annotation elements here (see also Figure 4a for examples) and provide full documentation of the annotation mechanism on LISTER’s GitHub repository (https://github.com/CPCLab/lister) and in the SI (chapter 1, with an illustration of the MM, provided in the SI Chapter 4).

a. *Basic elements*. A KV pair is represented as `{value|key}` in an experiment documentation entry. A KV pair can be extended with `measure` (denoting the measured quantity) and the `unit`, if necessary. Due to this extension, there are two more variations on how a KV pair may be written: `{measure|unit|key}` (in which the measure and unit will be mapped into value, with the unit at the end) and `{measure|unit|value|key}` (in which the measure and unit will be taken as given) (Figure 4a).
b. *Key visibility in .docx output*. In most cases, it is superfluous to have the *key* part of a KV pair available in the .docx output, as illustrated in Figure 4b for the key “expression media”. Hence, keys in the experiment documentation are hidden by default in the .docx output. However, if the key needs to be explicitly shown in the .docx output, users can indicate this by wrapping the key with colons “: …:”, i.e., `{value|:key:}` (Figure 4a).
c. *Order*. Each extracted KV pair is assigned an ‘order designator’ to disambiguate the order mapping of the keys. The order designator is derived from the paragraph number where the KV pair appears (Figure 4b).
d. *Comments*. There are three different types of comments supported in LISTER.

i. Comments that are parsed as-is, retaining both brackets and content in the .docx output. This is used to retain comments without modifications, both in the eLabFTW experiment documentation entry and in the .docx output. Such comments are marked as “`(this example)`”, which will be written as `(this example)` in the .docx output, whereas nothing is written in the metadata output (see SI chapter 1.3).
ii. Invisible comments with removed annotations. This is used when additional notes need to be specified in protocols or MMs that should be hidden from the .docx output. Typical examples are to detail 1) the meaning of a specific parameter, 2) the protocol/MM entry author(s), or 3) the protocol/MM entry version. Such comments are marked as “`(_invisible comment_)`”.
iii. Comments for which the content is retained but not the brackets. This is used for comments inside KV pairs. For example, “`the {empty vector strain (:as:)|:negative control:}`” will be written as “`the empty vector strain as negative control`” in the .docx output, with “`negative control`” as the key and “`empty vector strain`” as the value.
e. *Conditionals and iterations*. LISTER supports documenting conditionals and iterations in protocol and MM templates. Conditionals may be used when a step has multiple possibilities, with each possibility leading to a specific result, further steps to take, or termination. Iterations may be used when one or more steps need to be done repetitively until one or more specific condition(s) is (are) satisfied. See Table 1 for the three supported iteration types. Nonetheless, these elements should be used with caution in the final experiment documentation. When adapting the templates, researchers are encouraged to resolve conditionals by documenting the actual results or steps taken, thereby removing the alternatives from the experiment documentation. Likewise, for iterations, researchers are encouraged to document the respective repetitive steps explicitly.
f. *Reference management*. A reference in the LISTER annotation is supported by using a bracketed DOI (see SI, chapter 1.3). The annotation is parsed and listed as a numerical reference annotated in squared brackets in the text, with the referenced publications provided as a list of DOI at the end of the .docx document. DOIs are recognized based on their patterns using regular expressions.
g. *Sections*. The `<section | section name>` annotation is designated to provide a separation between sections.

**Table 1.**
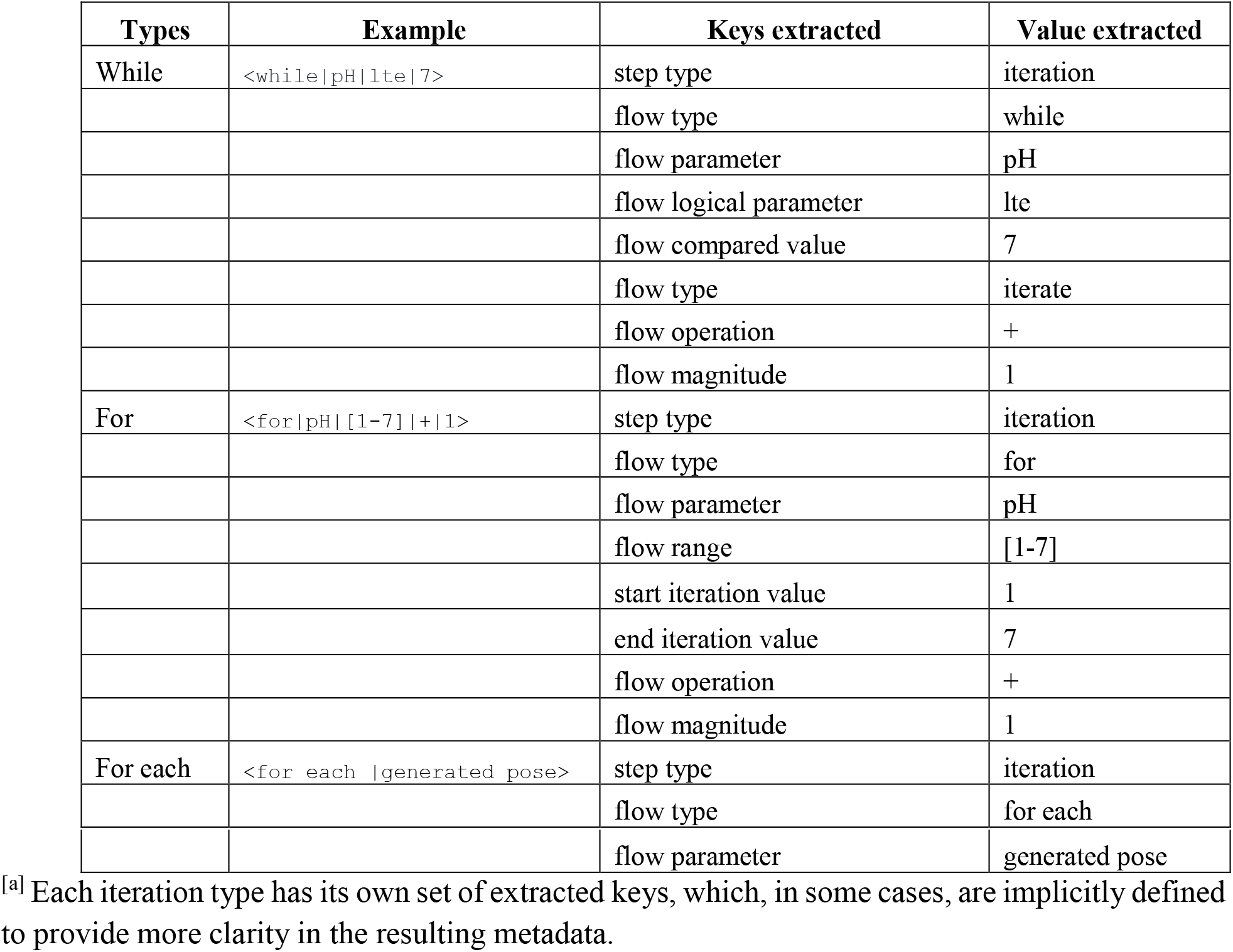
Three supported types of iteration, along with examples and extracted keys.

An entry of the *Experiment* type is parsed according to the above LISTER annotation rules. By contrast, *System, Project, Publication*, and *Study* entries are parsed for their attributes given in Figure 2, i.e., the table in such an entry is transformed into KV pairs.

#### Implementation of the LISTER app

LISTER is open-source under the GPLv3 license^16^ and implemented using Python 3.9. Metadata elements (key, value, measure, unit, comment, conditional/iteration, reference, section/subsection) are identified through regular expressions. We used the following Python libraries to develop LISTER: `BeautifulSoup`^17^ for parsing HTML content, `elabapy`^18^ and `elabapi-python`^19^ to communicate with the eLabFTW API endpoint, `json, xlsxwriter`^20^, and `python-docx`^21^ to write files in .json, .xlsx, and .docx formats, respectively. `Gooey`^22^, a Python library to create a GUI on top of a command line, is used to provide a graphical layer for LISTER. Other utility libraries used are `re` (regular expression), enum (enumeration), `os, PyInstaller` application packager^23^, `ssl, platform, pathlib, pandas`, and `lxml`^24^. For utilizing LISTER without having to configure Python and install Python libraries, LISTER has been packaged for different operating systems: Windows 10 and 11, Ubuntu-based Linux distributions (tested on Linux Mint 21), and macOS 10.12, 12.4, and 13.0. The code documentation is provided using the reStructuredText (reST) format^25^.

Parameters detailing the respective input source for syntax checking and metadata extraction are provided via the GUI (Figure 6). For experiment documentation coming from eLabFTW, the experiment ID (which is indicated on the URL of the experiment), API endpoint URL, and eLabFTW API Token are necessary. Both the API endpoint URL and eLabFTW API Token can be obtained from the administrator of the eLabFTW instance. The outputs are provided in the user-specified output directory. An accompanying ‘`config.json`’ file allows preloading parameters to the GUI, including the output file/directory names, and eLabFTW-specific parameters (e.g., experiment ID, API endpoint URL, and eLabFTW API Token) to ease the startup with the LISTER app or allow batch processing.

## EVALUATION

We introduced LISTER as a methodological and algorithmic solution for the field of life sciences to extract metadata from primary data using minimum efforts and making the primary data accessible and downloadable. LISTER consists of three components, customized eLabFTW entries, a container concept within eLabFTW, and the app. LISTER is available at https://github.com/CPCLab/lister.

As proof-of-concept and for our research documentation, we generated 11 annotated MM templates for the domain of computational biophysical chemistry and 4 templates for the domain of protein biochemistry and molecular biology. Generation of these templates took about 80 man-hours, including cross-correction rounds, accompanied by less than six meetings to define the granularity and scope of the MMs. These MM templates are being used in our research group to follow the designed RDM workflow. The MM templates are available at https://github.com/CPCLab/materials-and-methods to initiate the development of research group-specific templates. An illustrative example is provided in https://github.com/CPCLab/lister-container, containing a publication entry within eLabFTW and its corresponding system, project, study, and experiment, which is provided as an eLabFTW export containing CSV and PDF files. The extraction of the metadata took about 8 seconds for an experiment linked with corresponding entries (e.g., System, Project, and Study), and 20 seconds for a publication containing experiments with over thirty attachments in total, each linked with corresponding entries. We also set up a demo eLabFTW server for the user to try LISTER. The demo server is available at https://elabftw.pharm.hhu.de, and the config.json file is shared at https://github.com/CPCLab/lister, including the manual to reuse/modify the configuration file.

LISTER builds on the ISA Model and eLabFTW. The choice of the ISA Model reflects its support in the communities of life sciences, biomedicine, and environmental research^8^ and its straightforward and realistic structuring of heterogeneous experiments. Our extensions in terms of *Project, Publication*, and *Protocols/MM* (container) classes were motivated to map complex research environments, where more than one (funded) project addresses an overarching research question, summarize those experiments that have been entered into a publication, and store annotated protocols and MM sections to facilitate experiment documentation. The choice of eLabFTW as ELN reflects its design for the life sciences, which fits our user community, wide-spread use, active development, and facilitated access via the API, which provides us with experiment documentation and container class entries for the metadata parsing. An alternative ELN, Chemotion^26, 27^, is more focused on the chemistry domain. Note, though, that the ELN Consortium^28^, involving ELNs and RDM-related works including Chemotion, eLabFTW, Chemedata^29, 30^, Herbie^31^, Juliabase^32^, Kadi4Mat^33^, PASTA-ELN^34^, and SampleDB^35, 36^, aims at making ELN entries interchangeable via the ELN data format^28^.

We use DSpace^14^ to store, manage, and catalog primary data and metadata extracted by LISTER. DSpace has been specifically designed to provide storage, access, and preservation of digital archives on a long-term basis and is used by more than two thousand organizations worldwide^37^. Since DSpace is free and open-source software, it can be customized to create an adapted data preservation strategy. We run a customized DSpace instance that allows users to search the content of the contextual, LISTER-generated metadata along with the primary research data. At present, this instance is intended for research data storage and cataloging within the scope of our University but will be extended to long-term storage and (hierarchical) public access (fourth requirement mentioned above). Note that specifications of long-term storage may vary between labs and research environments because of differences in, e.g., storage size and data management practices. LISTER-extracted metadata is not limited to use in connection with our own DSpace-based archival but can be adapted for or uploaded to other (public) data repositories.

How does LISTER comply with the FAIR^6^ guidelines? In terms of *findability*, LISTER’s RDM workflow is able to create rich contextual metadata by using semi-automatic metadata extraction; the metadata can later be indexed to enable a search over the data cataloging platform DSpace. In terms of *accessibility*, the research data is published on the web using DSpace, and the HTTP/S protocol is used to access it. Regarding *interoperability*, metadata are serialized using the JSON format for data exchange. Metadata is also provided in .xlsx format for human readability. We do not yet use ontologies to represent the metadata in the Resource Description Framework (RDF)^38^ format, i.a., due to the limited availability of ontologies in our pilot domain computational biophysical chemistry. RDF allows information to be represented as a triple unit, consisting of the subject, predicate, and object. Each triple can be connected to other triples, yielding interconnected triples as linked data in the form of a knowledge graph, which can then be linked to another knowledge graph, making connected data well-integrated. We intend to incorporate alignment with ontologies or use the output from LISTER’s metadata extraction for collecting ontological terms in the future. Finally, concerning *reusability*, accurate and relevant attributes can be extracted using LISTER as it directly parses experiment documentation derived from annotated protocols/MMs and adapted by the researcher, which should minimize the number of inaccuracies or omissions.

How does LISTER relate to existing public data repository concepts? Several general-domain data repositories have been established for public use, e.g., Zenodo^39^, Dryad^40^, FigShare^41^, and Open Science Framework (OSF)^42^. A comparison of some of these repositories as to the storage quota on its free tier, possible storage extension and additional costs, DOI provision, funding source, the base of operations, and whether it is required to make contextual metadata available is given in Chapter 5 of the SI (SI Table 2). To make the published data more compliant with FAIR guidelines, key-value-based contextual metadata can be added. However, the contextual metadata of uploaded research data items is often not available, and when it is available, the content of contextual metadata, uploaded as an additional file(s), is not necessarily searchable in these portals. By contrast, the LISTER-based RDM workflow provides contextual metadata along with the primary data, and the customized DSpace instance allows searching it.

Complementary to the general-domain data repositories, there are also specific-domain data portals such as *Protein Data Bank* (*PDB*), *Universal Protein Database (UniProt*), and *Ensembl. PDB*^43^ is a well-established repository of resolved structures of biomolecules and their associated primary data as well as derived data. *UniProt*^4^ is a protein sequence database organized as UniProt Knowledge Base, Archive, and Reference Clusters. *Ensembl*^45^ is a database for genomic information. In these cases, the repositories’ content can be browsed by specific annotations^46–48^, which utilize vocabularies or ontologies organized by the Gene Ontology Consortium^49^ for categorization. Although it will be helpful to have a semantic annotation on LISTER-generated metadata, the semantic and ontological alignment is a wide topic and beyond the current work. The LISTER workflow lays the groundwork for semantic alignment by making contextual metadata available in the first place.

How does LISTER relate to other domain-specific RDM workflow concepts? The NFDI consortium DataPLANT^5, 50^ created technology stacks to manage research data in plant science, namely Annotated Research Context (ARC) and Swate. ARC provides a packaging mechanism for research data that includes measurement data, metadata, data annotations, tools, and scripts surrounding the research cycle on plant science. In addition to ARC, Swate^51^ is a plugin implemented for Microsoft Excel that allows annotating research using contextual metadata with alignment to specific and standardized ontologies^50^. Compared to either approach, LISTER does the packaging/archival of experiment documentation through the ELN and, thus, is more tailored and integrated toward ELN users. Furthermore, LISTER disentangles the creation of metadata and ontology alignment into two steps, with the first supported by protocol/MM templates and the second intended to be automated semantic metadata annotation via the use of external semantic terminology services, as provided, e.g., by NFDI4Chem^52^. We envision that this way, LISTER reduces the barrier of creating metadata for experiment documentation, albeit without immediate semantic annotation as a trade-off.

LISTER bears some similarity to the work of Schröder et al.^12^, which also uses eLabFTW to store experiment documentation. The authors provide a proof of concept in the Calcium imaging domain by implementing an automated semantic metadata extraction from manually engineered protocols with a canonical experiment entry structure. The protocols require domain expertise to write, also to ensure that the protocol/database elements are mapped to the correct vocabulary term or an ontological class instance. While this approach directly provides semantic annotation for protocols, it requires that domain ontologies for annotating the protocols already exist and that the protocol writer has the background to associate the correct ontological annotations to the protocols. These two requirements are not necessarily met in other scenarios, which resulted in the above-mentioned two-step approach by LISTER.

The Helmholtz Metadata Collaboration (HMC) focuses on facilitating research data documentation and handling across various fields for the Helmholtz Association. The HMC comprises several tools and services ranging from, among others, FAIR data publication guidelines, metadata specification/validation/structuring/sharing tools, and services to find metadata standards and provides persistent URLs^53^. LISTER, as a metadata extraction tool, could potentially be aligned with, for example, the HMC metadata specification and validation tool to validate the extracted metadata.

## CONCLUSION AND FUTURE WORK

We introduced LISTER as a methodological and algorithmic solution to extract metadata from primary data using minimum efforts and making the primary data accessible and downloadable. LISTER is tailored to the field of life sciences, makes use of existing RDM standards and platforms, and consists of three components: customized eLabFTW entries, a ‘container’ concept in eLabFTW, and an app to enable easy-to-use, semi-automated metadata extraction from eLabFTW entries. For our research documentation and as a showcase, we apply LISTER to the fields of computational biophysical chemistry as well as protein biochemistry and molecular biology. Our concept of reusable protocol and MM templates to derive documentation from experiments, along with extracting metadata from annotated experiment entries, should also be extendable to other life science areas.

Future work will aim at standardizing terms that occur in the extracted experiment metadata; such terms can then be aligned with existing ontologies or could also be used to extend related ontologies. We envision that deep learning-based approaches for the automatic labeling of keys or units in experiment documentation might become available when enough training data for a specific domain exists. LISTER may contribute to generating such training data.

## ASSOCIATED CONTENT

### Supporting Information

Source code and packaged apps for Windows, Linux, and macOS (both for x86-64 and arm64 architectures) are available at https://github.com/CPCLab/lister under the GPLv3.

An exemplary container class to adopt the LISTER workflow is available at https://github.com/CPCLab/lister-container, and a collection of MM templates for computational biophysical chemistry is available at https://github.com/CPCLab/materials-and-methods. Both the containers and MMs can be imported into a running eLabFTW instance.

## AUTHOR INFORMATION

### Author Contributions

H.G. conceptualization, supervision, analysis; F.M. conceptualization, investigation, programming, analysis, visualization; K.R. analysis, project management. The manuscript was written with the contributions of all authors. All authors have given approval for the final version of the manuscript.

## ACKNOWLEDGMENT

We are grateful to our colleagues who have contributed to writing the MM templates to be further processed by LISTER, Stephan Schott-Verdugo, Michele Bonus, Jesko Kaiser, Christoph Gertzen, Stefanie Brands, Jonas Dittrich, Christopher Pfleger, Christina Gohlke, and Filip König.

This work is guided by our experiences in the Collaborative Research Center (CRC) 1208. We thank Lutz Schmitt, Stefanie Weidtkamp-Peters, and other CRC1208 members for continued discussions. We are grateful for discussions with and support from the Research Data Management Competence Center (Dirk Fleischer, Rafael Dellen) and the Zentrum für Informations- und Medientechnologie (Nina Knipprath, Bert Zulauf, Sebastian Manten, and Thomas Dziurzyk) at Heinrich Heine University Düsseldorf.

## ABBREVIATIONS

ACS: American Chemical Society
API: Application Programming Interface
ARC: Annotated Research Context
DFG: Deutsche Forschungsgemeinschaft (German Research Foundation)
DOI: Digital Object Identifier
eLabFTW: electronic Lab For The World
ELN: Electronic Laboratory Notebook
FAIR: Findable, Accessible, Interoperable, and Reusable
GUI: Graphical User Interface
HMC: Helmholtz Metadata Collaboration
HTML: HyperText Markup Language
HTTP/S: Hypertext Transfer Protocol/Secure
ID: Identifier
ISA: Investigation-Study-Assay
JSON: JavaScript Object Notation
KV: Key-Value
LISTER: (LI)fe (S)cience Me(t)adata Pars(er)
MM: Materials and Methods
NFDI: Nationale Forschungsdateninfrastruktur (National Research Data Infrastructure)
NFDI4Chem: NFDI for Chemistry
NSF: National Science Foundation
OSF: Open Science Framework
PDB: Protein Data Bank
RDF: Resource Description Framework
RDM: Research Data Management
SOP: Standard Operating Procedures
UniProt: Universal Protein Database
URL: Uniform Resource Locator.

## DATA and SOFTWARE AVAILABILITY

The LISTER source code is available at https://github.com/CPCLab/lister. eLabFTW version 4.4.3 used here is available at https://www.elabftw.net/, the MM templates generated in this work are available at https://github.com/CPCLab/materials-and-methods and the class definitions for our eLabFTW adoption is available at https://github.com/CPCLab/lister, which can be imported to other eLabFTW instances as well.

