## Supporting Information for "LISTER: Semi-automatic metadata extraction from annotated experiment documentation in eLabFTW"

The following is a detailed description of technical details for LISTER, as it is also available in the top-level README file in the LISTER repository. For the most current version of this description, see https://github.com/CPCLab/lister.

### 1. LISTER: Life Science Experiment Metadata Parser

This repository contains a set of files to parse documentation of experiments in eLabFTW.

#### 1.1 Motivation

As a research group usually has its own set of experiment protocols/Materials and Methods (MM) to conduct experiments, a tool to extract metadata from protocols/MMs-adapted experiment documentation helps support research data publication according to FAIR (Findable, Accessible, Interoperable, and Reusable) principles. Research data published under FAIR principles is expected to improve the reproducibility of research. To enable metadata extraction, the experiment documentation follows annotation rules described below and is in an HTML format stored as an experiment/database entry in eLabFTW.

#### 1.2 Installing and running LISTER

LISTER is distributed as an executable file for Windows, Linux, or macOS (with an Intel chipset). The executable file for each platform is available on the release page, along with another, platform-specific file.

- **For Windows and Linux**, place the executable file (`lister.exe` on Windows or `lister` on Linux) within the same folder as the `config.json` file.
- **For macOS**, create the directory `~/Apps/lister` first and place the executable `lister.app` and `config.json` in this directory.

##### 1.2.1 Adapting the config.json file

Parsing an eLabFTW entry requires

- the general parameters
  - eLabFTW `API token` and `API endpoint`, which can be obtained from the eLabFTW instance's administrator of the lab or university,
  - Default `output` directory, i.e., a directory path used to store the parser output,

and

- experiment-specific parameters

Metadata/experiment output filename and Experiment ID or Database ID for the entry to be parsed.

#### 1.3 Annotation mechanism

The annotation mechanism allows extracting metadata from experiment documentation as .xlsx and .json files. In the following points, the basic elements of annotating protocol/MM to be parsed by LISTER are described.

- *Key-Value (KV) elements*.
    - A KV pair is written as `{value|key}` in an experiment entry.
    - If applicable, a KV pair is extendable with `measure` and `unit`. Therefore, there are two more variations for writing a KV pair:
        - `{measure|unit|key}` the measure and unit will be mapped into value and unit.
        - `{measure|unit|value|key}` the measure and unit will be taken as given.
    - For example, "*Two* *`{100|mL|LB Kan|expression media}`* *`cultures in {unbaffled Erlenmeyer|flasks}`*" consists of two patterns of pair:
        - `{measure|unit|value|key}` → *`{100|mL|LB Kan|expression media}`*
        - *`{value|key}`* → *`{unbaffled Erlenmeyer|:flasks:}`*"
    - Keys are hidden by default in the .docx output file to avoid superfluous text.
    - To make the keys visible, they can be placed within colons as `{value|:key:}`, such as *`{unbaffled Erlenmeyer|:flasks:}`*".
- *Order*.
    - As there can be identical keys within an experiment entry, disambiguation is needed.
    - The disambiguation is done through the *paragraph number*, which will be extracted and associated with each KV pair.
- *Comments*. There are three types of comments supported in LISTER.
    - Comments parsed as-is.
        - This retains both brackets and content in the word document.
        - Annotation is done using a regular bracket ().
        - Annotation example: `(This comment will be parsed as is, retaining both the content and the brackets in the .docx file.)`.
    - Invisible comments.
        - Used to specify additional notes (regarding, e.g., parameter use) that should be hidden from the final experiment documentation output.
        - Annotation is done using a pair of underscores inside a regular comment. (_ _)
        - Annotation example: `(_This comment will be invisible in the .docx output file._)`.
    - Comments are retained but without brackets.

        - This is typically used for comments within KV pairs.
        - Annotation is done using brackets and a double colon (: :)
        - Annotation example: `(:This comment's bracket will be invisible in the .docx output file, but the text content will be kept.:)`.

- *Conditionals and iterations handling.*
    - LISTER supports documenting conditionals and iterations, but this should be used cautiously: As the final experiment documentation is unlikely to have these conditional and iteration clauses, researchers are required to resolve them by adapting the experiment parameter values with the actual values used during the experiment.
    - Annotation example:

```
<if|monomer|e|true>
   With {monomer|:structure:}, the target sequence was provided in
form of fasta file and uploaded to the server.
<elif|multimer|e|true>
   With {multimer|:structure:}, the fasta file with the {3| number
   of chains in the target complex} target sequences were prepared
   and uploaded to the server.
</if>
```

    - The users are expected to adopt the conditionals after importing the MM into the experiment, by removing the non-relevant clause and also removing the header of the relevant clause. For example, if one uses only the multimer, this would be the remaining MM lines in the experiment:

```
With {multimer|:structure:}, a fasta file with the {3| number of
chains in the target complex} target sequences were prepared and
uploaded to the server.
```

- *Reference management.*
    - References can be provided if the referred source has a DOI.
    - Annotation is done using regular brackets and providing the DOI (not URI) in the bracket.
    - The DOI will be converted into Arabic numerals in square brackets, which refer to the reference provided at the bottom of the document.
    - References are only retained in the docx output, but not the metadata outputs (.xlsx/.json).
    - Example: `(DOI_CODE)`, such as `(10.1073/pnas.062492699, 10.1002/elsc.200800043)` will be written as `[1]` in the experiment body, and as a numbered list of DOI codes at the end of the experiment documentation.
- *Sections.*
    - The keywords `section` or `subsection` are designated to provide a separation between sections or subsections.
    - This is done by using the `<section|section name>` annotation.
    - Multiple subsections are also supported with `<subsubsection|section name>`, which will output different sectioning levels in the .xlsx and .json files and different heading levels in the .docx file.

#### 1.4 Examples of annotations vs. extracted metadata

<table>
<tr><th>Extracted item</th><th>Description</th><th>Representation</th><th>Example</th><th>Extracted order, key, value, and optionally measure, unit in the metadata</th></tr>
<tr><td>Section</td><td>The section name</td><td><section|section name></td><td><section|Structure Preparation></td><td><ul><li>"-", section level 0, Structure Preparation, -, -</li></ul></td></tr>
<tr><td>Order</td><td>The <code>order</code> of the steps, based on the order of the paragraph in the experiment documentation</td><td>-</td><td>-</td><td>-</td></tr>
<tr><td>Key</td><td>The key for the metadata, connected to the value {value|key}.</td><td>{value|key}</td><td>{sequence alignment|stage}</td><td><ul><li><order>, stage, sequence alignment, -, -</li></ul></td></tr>
<tr><td>Comment, please also see the bullet points about comments above for variations</td><td>Comments are allowed within the key-value annotation, represented within regular brackets. Comments can be placed both/either before and/or after the key and/or value.</td><td>{value|(comment) key} or {value (comment)|key} or {value (comment)|(comment) key}</td><td>{receptor residue|(minimization) target}</td><td><ul><li><order>, target, receptor residue, -, -</li></ul></td></tr>
<tr><td>Value</td><td>The value of the metadata is the first item within the curly brackets {value|key}.</td><td>{value|key}</td><td>{sequence alignment|stage}</td><td><ul><li><order>, stage, sequence alignment, -, -</li></ul></td></tr>
<tr><td>Measure and Unit</td><td>The measure and unit of corresponding key/value pairs.</td><td>{measure|unit|value|key}</td><td>{100|mL|LB Kan|expression media}</td><td><ul><li><order>, expression media, LB Kan, 100, mL</li></ul></td></tr>
<tr><td>Value and Unit</td><td>In some cases, value is attached to a unit directly, without having to provide a measure.</td><td>{value|unit|key}</td><td>{250|rpm|shaking}</td><td><ul><li><order>, shaking, 250, -, rpm</li></ul></td></tr>
</table>

| Extracted item | Description | Representation | Example | Extracted order, key, value, and optionally measure, unit in the metadata |
|---|---|---|---|---|
| Control flow: `for each` | Extract multiple key-value pairs related to `for each` iteration | <for each\|iterated value> | <for each\|*generated pose*> | ▪ <order>, step type, *iteration*, -, -<br>▪ <order>, flow type, *for each*, -, -<br>▪ <order>, flow parameter, generated pose, -, - |
| Control flow: `for` | Extract multiple key-value pairs related to `for` iteration | <for\|key\|[range]\|iteration operation\|magnitude> | <for\|pH\|[1-7]\|+\|1> | ▪ <order>, step type, *iteration*, -, -<br>▪ <order>, flow type, *for*, -, -<br>▪ <order>, flow parameter, pH, -, -<br>▪ <order>, flow range, [1-7], -, -<br>▪ <order>, start iteration value,1, -, -<br>▪ <order>, end iteration value,7, -, -<br>▪ <order>, flow operation, +, -, -<br>▪ <order>, flow magnitude, 1, -, - |
| Control flow: `while` | Extract multiple key-value pairs related to `while` iteration | <while\|key\|logical operator\|value> ... <iterate\|iteration operation\|magnitude> | <while\|pH\|lte\|7> ... <iterate\|+\|1> | ▪ <order>, step type, *iteration*, -, -<br>▪ <order>, flow type, *while*, -, -<br>▪ <order>, flow parameter, pH, -, -<br>▪ <order>, flow logical parameter, lte, -, -<br>▪ <order>, flow compared value, 7, -, - |

<table>
<tr><th>Extracted item</th><th>Description</th><th>Representation</th><th>Example</th><th>Extracted order, key, value, and optionally measure, unit in the metadata</th></tr>
<tr><td></td><td></td><td></td><td></td><td>▪ <order>, flow type, iterate (after while)<br>▪ <order>flow operation, +, -, -<br>▪ <order>, flow magnitude, 1, -, -</td></tr>
<tr><td>Control flow: <code>if</code></td><td>Extract multiple key-value pairs related to <code>if</code> iteration</td><td><code><if|key|logical operator|value></code></td><td><code><if|pH|lte|7></code></td><td>▪ <order>, step type, conditional, -, -<br>▪ <order>, flow type, if, -, -<br>▪ <order>, flow parameter, pH<br>▪ <order>, flow logical parameter, lte, -, -<br>▪ <order>, flow compared value, 7</td></tr>
<tr><td>Control flow: <code>else if</code></td><td>Extract multiple key-value pairs related to else if iteration</td><td><code><else if|key|logical operator|value></code></td><td><code><else if|pH|between|[8-12]></code></td><td>▪ <order>, step type, conditional, -, -<br>▪ <order>, flow type, else if, -, -<br>▪ <order>, flow parameter, pH, -, -<br>▪ <order>, flow logical parameter, between, -, -<br>▪ <order>, flow range, [8-12], -, -<br>▪ <order>, start iteration value,8, -, -<br>▪ <order>, end iteration value,12, -, -</td></tr>
</table>

| Extracted item | Description | Representation | Example | Extracted order, key, value, and optionally measure, unit in the metadata |
|---|---|---|---|---|
| Control flow: `else` | Extract multiple key-value pairs related to else iteration | `<else>` | | ▪ `<order>, step type, *conditional*, -, -`<br>▪ `<order>, flow type, *else*, -, -` |

##### 1.4.1 Supported operators

###### *1.4.1.1 Logical operator*

A logical operator is used to decide whether a particular condition is met in an iteration/conditional block. It is available for while, if, and else if control flows. The following logical operators are supported:

- `e`: equal
- `ne`: not equal
- `lt`: less than
- `lte`: less than equal
- `gt`: greater than
- `gte`: greater than equal
- `between`: between

###### *1.4.1.2 Iteration operator*

An iteration operator is used to change the value of a variable in a loop. It is available for `while` and `for`. The following iteration operators are supported:

- `+`: iteration using addition
- `-`: iteration using subtraction
- `%`: iteration using modulo
- `*`: iteration using multiplication
- `/`: iteration using division

#### 1.5 Document validation

LISTER checks and reports the following syntax issues upon parsing:

- Orphaned brackets.
- Mismatched data types for conditionals and iterations.
- Mismatched argument numbers for conditionals and iterations.
- Invalid control flows.

#### 1.6 Image extraction

Images are extracted from the experiment documentation, but there is no metadata or naming scheme for the extracted images.

#### 1.7 Recommendations

- Avoid referring to, e.g., a section without explicitly using a key-value pair (avoid, e.g., "*Repeat step 1 with similar parameters*"), as this will make the metadata extraction for that particular implicit step impossible.
- To minimize confusion regarding units of measurement (e.g., `fs` vs `ps`), please explicitly state the units within the value portion of the key-value pair, e.g., `{0.01|ps|gamma_ln}`.

#### 1.8 GitHub repository structure

- The base directory contains the metadata extraction script.
- The `output` directory contains the extracted metadata: step order – key – value – measure – unit in JSON and XLSX format.

#### 1.9 Miscellaneous

##### 1.9.1 Packaging LISTER

- Packaging is done through the `PyInstaller` library and has to be done on the respective platform. `PyInstaller` should be installed first.
- It is recommended to use virtual environments using python's `venv` or anaconda.
    - Using `venv`
        - create `venv` virtual environment inside lister directory named `venv`, which will use python3.9 as the interpreter: `python3.9 -m venv venv`
        - set IDE to use the created `venv` environment as python interpreter; in pycharm, it is in the *Settings* - *Project: lister* - *Python Interpreter* - *Add Interpreter*, which is set to `/lister/venv/bin/python`
        - activate the venv environment: `source venv/bin/activate`
        - install required libraries: `pip install xlsxwriter gooey python-docx elabapy beautifulsoup4 pyinstaller pandas latex2mathml`
        - package LISTER app for relevant operating systems using the build scripts mentioned below
- A `.spec` file to build LISTER can be generated using the `pyi-makespec` command, e.g., `pyi-makespec --onedir lister.py` to create a spec file to package the LISTER app as one directory instead of one file.
- The spec file for each platform is provided in the `build-scripts` folder of the LISTER GitHub repository.

- The resulting packaged app will be available under the `dist` directory, which is created automatically during the build process.

###### *1.9.1.1 Packaging the app on Windows*

- One directory version - on the root folder of the repo, run `pyinstaller .\build-scripts\build-windows-onedir.spec`
- One file version - on the root folder of the repo, run `pyinstaller .\build-scripts\build-windows-onefile.spec`

###### *1.9.1.2 Packaging the app on Linux*

- One file version - on the root folder of the repo, run `pyinstaller build-scripts\build-linux-onefile.spec`

###### *1.9.1.3 Packaging the app on macOS*

- One file version - on the root folder of the repo, run `pyinstaller build-scripts\build-macos-onefile.spec`

#### 1.10 Troubleshooting

##### 1.10.1 Slow app execution

Decompressing a single-executable lister app into a temporary directory likely caused this problem. The multi-file distribution (aka one-directory version) can be used instead, although it is not as tidy as compared to the single-executable LISTER app.

##### 1.10.2 Windows: Encoding problem

When the following error `'charmap' codec can't encode characters in position...` appears, open `cmd.exe` as an administrator before running LISTER and type the following:

```
setx /m PYTHONUTF8 1
setx PATHEXT "%PATHEXT%;.PY"
```

##### 1.10.3 Windows: Packaging failure

The error `win32ctypes.pywin32.pywintypes.error: (110, 'EndUpdateResourceW', 'The system cannot open the device or file specified.'` happens because of file access problems on Windows. Check if the directory is not read-only, exclude the repo folder from antivirus scanning, and/or try removing both the `build` and `dist` directories. Both of these directories are automatically generated upon packaging. Cloud storage synchronization may also be the cause of this issue.

##### 1.10.4 macOS: dependencies not included

Please consider using an environment management system such as anaconda to package the app. Install Conda locally along with the dependencies stated in the `requirements.txt`. In the release, python 3.9.15 was used. LISTER runs fine on macOS v13.0.1 and macOS v10.12.4 within intel machines.

##### 1.10.5 macOS: unable to get into GUI

Running `lister.py` directly from your IDE on macOS may lead to the following message:

```
This program needs access to the screen. Please run with a Framework build of python, and only when you are logged in on the main display of your Mac.
```

Run the script from the terminal using `pythonw lister.py` instead.

### 2. Attributes for System, Project, Study, and Publication entry types

**Table S1.** Customized classes created for the eLabFTW database along with their attributes.[1]

| (a) System | (b) Project | (c) Study | (d) Publication |
|---|---|---|---|
| System title | Project title | Study aim | Publication title |
| Responsible person for the system | Project subtitle | Study responsible person | Publication authors |
| Further information | Project type | Study starting date | Published in journal |
| | Project cooperation partners | Study end date | Publication status |
| | Project manager | Further information for the study | DOI |
| | Project responsible person in the working group | | |
| | Project staff | | |
| | Project starting date | | |
| | Project end date | | |
| | Project status | | |
| | Project further information | | |

[1] (a) System - provides a general overview of the studied field. (b) Project - describes the specific project – type, funding information, people involved, and start/end date. (c) Study - provides the study aim, involved people, research data location, and publication list. (d) Publication –defines particular publication information and links the publication with experiments conducted as well as the related system, project, and study.

### 3. Writing principles for protocols and MMs

For protocols and MMs to be broadly reusable as templates for experiment documentation, the following guidelines are suggested:

- **Sharing of domain expertise.** Domain experts decide what is deemed important to document for the experiment, and in particular, the key-value pairs that are relevant for the metadata. In the current LISTER implementation, we do not restrict what can be written as metadata and the data types of the written keys. Once a pattern regarding recurring keys and their data types has been established for a studied domain, restrictions can be introduced along with possible existing hierarchies (ontologies) among the extracted keys.

- **Granularity.** It is recommended to write the protocols/MMs granularly and specifically in terms of information encoded in the KV pairs to avoid ambiguous use of keys and values. For example, if applicable, writing KV elements as key, measure, and unit is preferable to key and value only.

- **Ambiguity prevention**. Ambiguity occurs when identical keys appear in the same paragraph, as both keys are assigned to the same order. Thus, splitting the paragraph is recommended such that identical keys appear in different paragraphs.

### 4. Structuring and annotating MMs in eLabFTW

As mentioned in the main text, an MM catalog is categorized as ‘MM’ in the eLabFTW database, with relevant tags provided for filtering with the eLabFTW filter functionality (Figure S1). The content of an exemplary MM is shown in Figure S2.

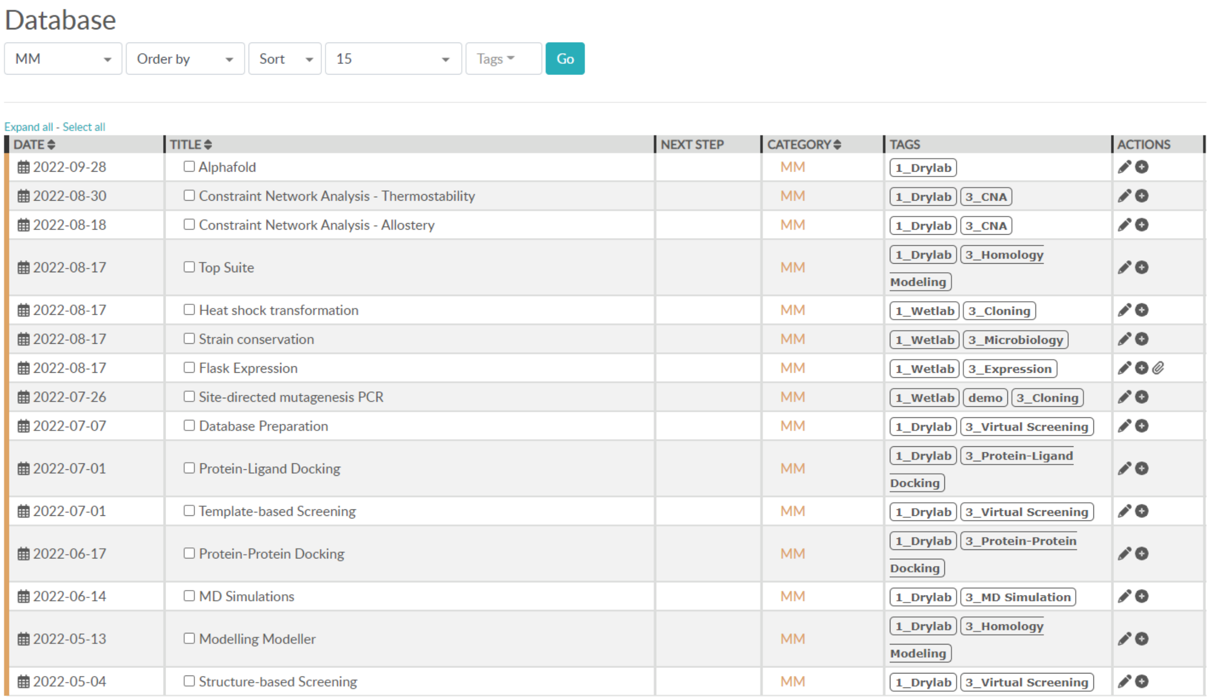

| DATE | TITLE | NEXT STEP | CATEGORY | TAGS | ACTIONS |
|---|---|---|---|---|---|
| 2022-09-28 | Alphafold | | MM | 1_Drylab | |
| 2022-08-30 | Constraint Network Analysis - Thermostability | | MM | 1_Drylab 3_CNA | |
| 2022-08-18 | Constraint Network Analysis - Allostery | | MM | 1_Drylab 3_CNA | |
| 2022-08-17 | Top Suite | | MM | 1_Drylab 3_Homology Modeling | |
| 2022-08-17 | Heat shock transformation | | MM | 1_Wetlab 3_Cloning | |
| 2022-08-17 | Strain conservation | | MM | 1_Wetlab 3_Microbiology | |
| 2022-08-17 | Flask Expression | | MM | 1_Wetlab 3_Expression | |
| 2022-07-26 | Site-directed mutagenesis PCR | | MM | 1_Wetlab demo 3_Cloning | |
| 2022-07-07 | Database Preparation | | MM | 1_Drylab 3_Virtual Screening | |
| 2022-07-01 | Protein-Ligand Docking | | MM | 1_Drylab 3_Protein-Ligand Docking | |
| 2022-07-01 | Template-based Screening | | MM | 1_Drylab 3_Virtual Screening | |
| 2022-06-17 | Protein-Protein Docking | | MM | 1_Drylab 3_Protein-Protein Docking | |
| 2022-06-14 | MD Simulations | | MM | 1_Drylab 3_MD Simulation | |
| 2022-05-13 | Modelling Modeller | | MM | 1_Drylab 3_Homology Modeling | |
| 2022-05-04 | Structure-based Screening | | MM | 1_Drylab 3_Virtual Screening | |

**Figure S1.** An exemplary list of MM entries together with associated tags.

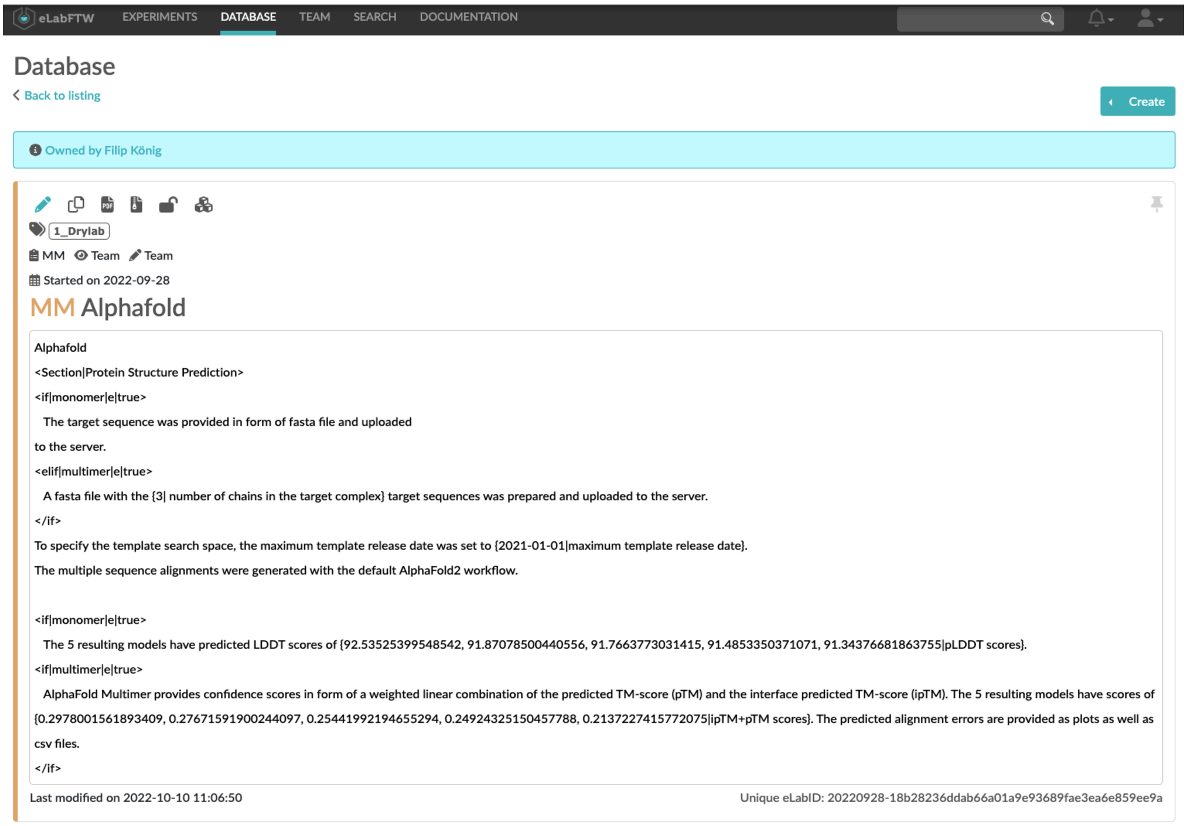

**Figure S2.** Exemplary content of an MM entry in eLabFTW with annotation markups that conform to LISTER's annotation rules. When a user imports this MM entry to their experiment, the content will be adapted according to the experiment.

### 5. Public Research Data Repositories

**Table 2.** A comparison of features across different research data repositories.

| Data repository | Zenodo | Dryad | FigShare | OSF |
|---|---|---|---|---|
| **Capacity free of charge** | 50 GB | - | 20 GB | 50 GB for the public, 5 GB for private |
| **Possible expansion** | Via contacting Zenodo's support; donations encouraged | 300 GB, extensible via contacting Dryad support; coverage fee for submissions outside of member institutions/journal | Via paid trier (Figshare plus), charged per dataset over a specific one-time Data Publishing Charge (DPC) / publication | External cloud services are supported, but not larger than OSF-provided storage |
| **Additional cost** | No | US$ 120 of base Data Publishing Cost, except for sponsor institutions/publishers or institutions from lower/mid-lower income nationalities. For non-sponsor/waived submission, data over 50 GB is charged for US$ 10 / extra 10GB. | Depending on the size | No |
| **DOI provision** | Yes, with versioning | Yes | Yes | Yes |
| **Funding** | CERN, EU-H2020, donations | Institution/publisher sponsorships, data upload fees | Private investors, such as Digital Science and Holtzbrinck Digital | Center for Open Science (Laura and John Arnold Foundation) - reserve funding of US$ 250.000 that would last 50 years, donations. |
| **Base of operation** | CH | US | UK | US |
| **The requirement to make contextual metadata available** | No | No | No | No |
